# Subcellular spatial proteomics of the heterotrophic nanoflagellate, *Cafeteria roenbergensis*

**DOI:** 10.64898/2026.10.13.758971

**Authors:** Dagmar Jirsová, Yu-Ping Poh, Katarzyna Dabrowska, Ritin Sharma, Patrick Pirrotte, Jeremy G. Wideman

## Abstract

*Cafeteria roenbergensis* is the best-studied free-living heterotrophic nanoflagellate, an abundant marine bacterivore and the host of the giant virus CroV, yet almost nothing is known about how its cell is organized. To provide broad insight into the localization of its proteome, we used subcellular spatial proteomics (SSP), based on a label-free version of LOPIT-DC (Localisation of Organelle Proteins by Isotope Tagging after Differential ultraCentrifugation), in which organelles are separated by differential centrifugation and each protein is assigned to a compartment from its abundance profile across fractions. We detected 5,948 proteins (5,889 protein groups), corresponding to 69.7% of the predicted *C. roenbergensis* proteome, and assigned high-confidence localizations to 2,945 proteins across 14 subcellular compartments, including basal body/cilia, cytosol, ER, Golgi, lysosome, membrane-trafficking, mitochondria, nucleoplasm, nucleus, peroxisome, plasma membrane, proteasome, ribosome, and spliceosome. This study serves as a proteomic dataset to resolve protein compartmentalization in a free-living heterotrophic stramenopile, enabling evolutionary, biological and functional insight and analysis. Investigators interested in using these data prior to publication are encouraged to contact us to avoid overlap and to facilitate coordinated and collaborative use of this resource.

## Introduction

Heterotrophic flagellated protists are abundant and ecologically important drivers of carbon turnover, yet their fundamental cell biology remains poorly understood. As major consumers of bacteria, these organisms form an essential link in aquatic food webs and drive nutrient cycling (Sanders et al., 1989; Hahn and Höfle, 2001; Jürgens and Massana, 2008). Thus, heterotrophic flagellates have been investigated primarily through the lens of taxonomy and ecology, leaving their cell biology comparatively unexplored. Heterotrophic flagellated stramenopiles like *Cafeteria* spp. are among the most widespread and offer an excellent system to investigate.

Stramenopiles (aka heterokonts) are protists characterized by their unequal flagella, one of which is tinseled with proteinaceous mastigonemes (Patterson, 1989). Along with two other major protist lineages, the alveolates (including ciliates, dinoflagellates, and apicomplexans) and the rhizarians (including cercozoans, foraminifera, and radiolarians), stramenopiles cluster into the billion-year-old supergroup SAR (Burki et al., 2007; Burki et al., 2020). Stramenopiles comprise two strongly supported clades, the Bigyra and the Gyrista (Cavalier-Smith and Chao, 2006; Riisberg et al., 2009; Derelle et al., 2016). Gyrista includes the predominantly osmotrophic and parasitic pseudofungi (e.g., oomycetes) and the only photosynthetic stramenopile clade, the ochrophytes (e.g., diatoms and kelp). Bigyra comprise several strictly heterotrophic lineages, including Bicosoecida, Labyrinthulomycota, and Opalinata, the latter of which includes the parasitic/commensal genus *Blastocystis* (Silberman et al., 1996; Moriya et al., 2000; Moriya et al., 2002; Derelle et al., 2016). Heterotrophy is ancestral in stramenopiles, so free-living heterotrophic flagellates retain the cellular toolkit from which the photosynthetic ochrophytes and osmotrophic pseudofungi were derived (Jirsová and Wideman, 2024). Though the distribution and ecology of some heterotrophic stramenopiles have been investigated, very little is known about the cell biology of the free-living heterotrophic flagellates within this group.

Even the best-studied heterotrophic flagellated stramenopile, *Cafeteria roenbergensis*, is poorly understood at the cell-biological level. *Cafeteria* spp. are among the most common and abundant components of marine plankton (de Vargas et al., 2015; Schoenle et al., 2020). *C. roenbergensis* cells are relatively small (∼8–10 µm) with two flagella, the anterior of which is equipped with mastigonemes (Fenchel and Patterson, 1988). Transmission electron microscopy of serial cross-sections has revealed a cytostome, a single nucleus, and up to five mitochondria per cell (O’Kelly and Patterson, 1996). *C. roenbergensis* has served as a model for studying bacterial grazing by nanoflagellates (Boenigk and Arndt, 2000; Jürgens and Massana, 2008) and has enabled characterization of the giant marine DNA virus CroV (Massana et al., 2007; Fischer et al., 2010). Despite more than three decades of study since its description (Fenchel and Patterson, 1988), high-quality genomes have only recently become available for four strains of *C. roenbergensis* (Hackl et al., 2020), opening the door to cell biological investigation through subcellular spatial proteomics (SSP).

SSP maps protein localization across the whole cell. Cells are gently lysed, organelles and other structures are separated into a series of fractions by centrifugation, and the abundance of each protein across the fractions is measured by quantitative mass spectrometry. Proteins that reside in the same compartment share a fractionation profile and can be assigned to that compartment by comparison with marker proteins of known location (Dunkley et al., 2004; Foster et al., 2006; Christoforou et al., 2016; Geladaki et al., 2019; Lundberg and Borner, 2019; Christopher et al., 2021; Breckels et al., 2024). We use a label-free version of LOPIT-DC (Localisation of Organelle Proteins by Isotope Tagging after Differential ultraCentrifugation), in which fractions are generated by differential centrifugation (Geladaki et al., 2019). First developed in plant and animal systems (Dunkley et al., 2006; Foster et al., 2006; Christoforou et al., 2016; Itzhak et al., 2017), SSP has since been applied across eukaryotic diversity, including budding yeast (Nightingale et al., 2019), apicomplexans (Barylyuk et al., 2020; Guérin et al., 2023; Chisholm et al., 2026), kinetoplastids (Moloney et al., 2023; Hammond et al., 2026), diplonemids (Hammond et al., 2025), ciliates (Jirsová et al., 2025; Chen et al., 2025), and the metamonad *Paratrimastix pyriformis* (Zítek et al., 2022). Here, we apply SSP to *Cafeteria roenbergensis* and localize 2,945 proteins to 14 cellular compartments, including basal body/cilia, cytosol, ER, Golgi, lysosome, membrane-trafficking, mitochondria, nucleoplasm, nucleus, peroxisome, plasma membrane, proteasome, ribosome, and spliceosome, providing the first cell-wide map of protein localization in a free-living heterotrophic stramenopile.

## Results and Discussion

*Cafeteria roenbergensis* RCC4623 was grown for 2–3 days at room temperature in f/2 medium supplemented with tryptic soy broth, until the culture reached stationary phase. The total culture volume was 4 L. Cells were harvested by centrifugation, washed twice with 1× PBS, and centrifuged after each wash. The final pellet was resuspended in DF buffer. The final volume used for cell lysis was 8 mL, containing approximately 14 × 10^6^ cells. Cells were gently lysed by nitrogen cavitation or passage through a filter (5 µm and 1.2 µm). These conditions preserved membrane-bound organelles, and the resulting homogenate was separated into 11 fractions by differential centrifugation following the LOPIT-DC protocol (Geladaki et al., 2019). The pellet from each centrifugation step, together with the supernatant remaining after the final spin, was collected for mass spectrometry analysis. The experiment was performed using the three different cell lysis methods. Protein recovery across the fractionation series was assessed by SDS-PAGE followed by Coomassie staining (Figure S1).

Label-free mass spectrometry analysis of the 11 fractions identified 5,948 proteins belonging to 5,889 protein groups, corresponding to 69.7% of the predicted *C. roenbergensis* proteome. Protein abundances were normalized within each experimental replicate, and missing values were imputed by neighbour averaging, with or without subsequent zeroing. Abundance profiles across the fractionation series were generated for each protein (Figure 1A) and visualized using t-distributed stochastic neighbor embedding (t-SNE) (van der Maaten and Hinton, 2008) to reveal protein clusters (Figure 1B).

**Figure 1.**
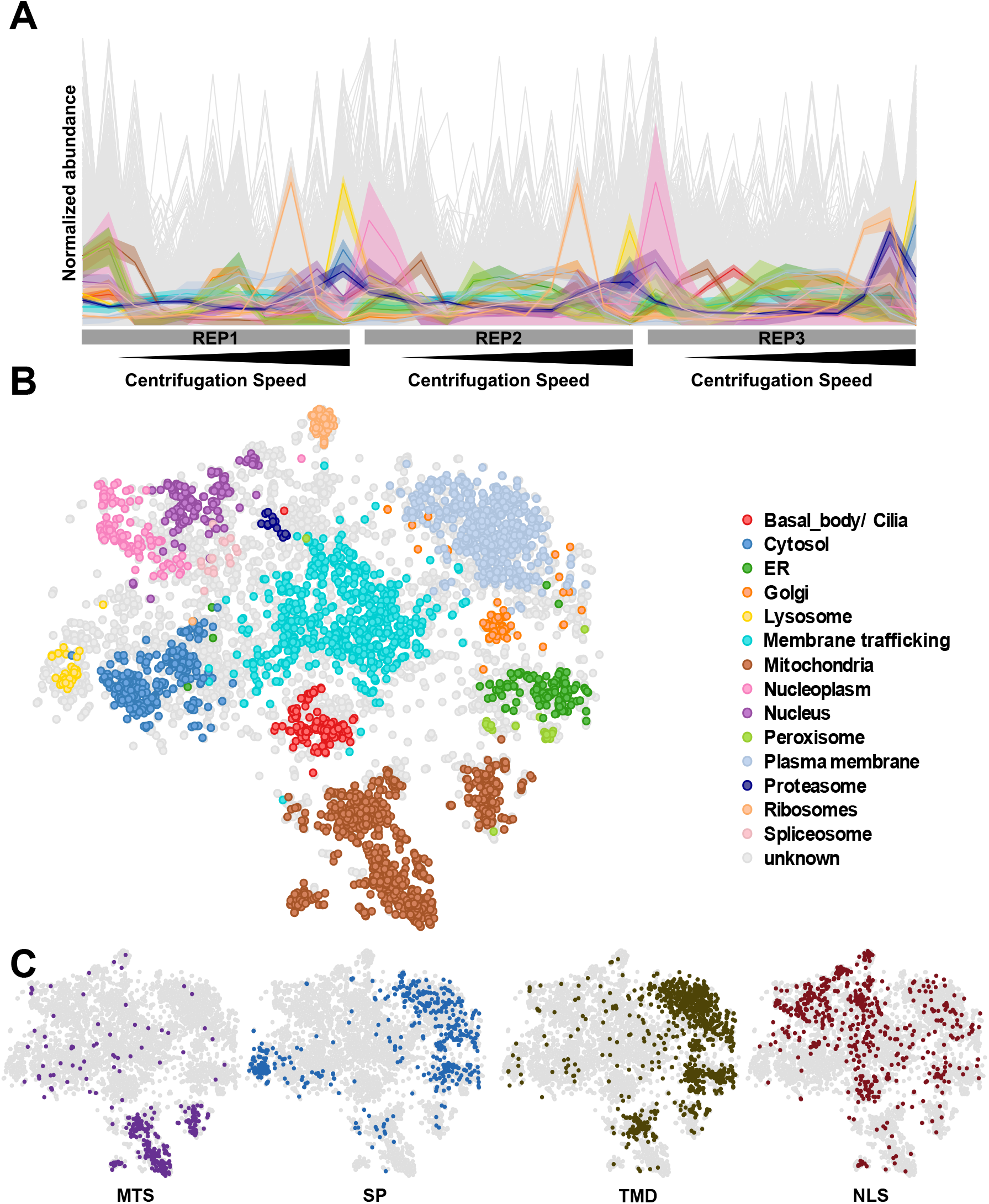
Subcellular proteome of *Cafeteria roenbergensis*. A. Normalized abundance profiles across the fractionation series for each of three independent experiments (REP1–REP3), with marker proteins coloured by compartment and all remaining proteins in grey. B. Protein localizations predicted by a supervised support vector machine classifier trained on 571 marker proteins across 14 compartments and filtered at a median probability cuto?, represented by t-SNE. C. Sequence-based targeting predictions mapped onto the same embedding, showing mitochondrial targeting signals (MTSs), signal peptides (SPs), transmembrane domains (TMDs) and nuclear localization signals (NLSs).

### Machine Learning-Based Assignment of Subcellular Localization

To evaluate the bias introduced by imputation, three datasets representing different levels of imputation were each classified by SVM using a manually curated marker set (Table S1). SVM assignments were consistent across the three datasets (Figure S2), and the dataset imputed by neighbour averaging followed by zeroing was therefore selected for downstream analysis in order to maximize protein coverage. Applying a median probability cutoff, the trained SVM assigned subcellular localizations to 2,945 *C. roenbergensis* proteins across 14 compartments (Figure 1B and Table S2), providing the first predicted localization for thousands of previously uncharacterized stramenopile proteins.

To assess the reliability of the SVM assignments, we compared them against sequence-based prediction tools: TargetP (mitochondrial targeting signals, MTS, and signal peptides, SP), DeepTMHMM (transmembrane domains) and NLStradamus (nuclear localization signals, NLS) (Figure 1C). The predicted mitochondrial proteins formed a well-defined cluster at the bottom of the t-SNE, closely mirroring the distribution of proteins carrying an MTS. Proteins with signal peptides and transmembrane domains were concentrated in the lysosome, plasma membrane, ER, and Golgi clusters, as expected for compartments of the secretory pathway, whereas NLS-containing proteins were enriched across the nucleus and ribosomal clusters. Overall, these sequence-level features align with the localizations predicted from the fractionation data. For comparison, unsupervised k-means clustering with k = 14, matching the number of marker-defined compartments, yielded somewhat similar clustering patterns, though some organelles were split into multiple clusters (e.g., mitochondria and plasma membrane), and others were merged with others (e.g., peroxisome) (Figure S2).

## Conclusions

Here we report the first subcellular proteome of *Cafeteria roenbergensis*, and the first for any free-living heterotrophic stramenopile. Heterotrophy is ancestral in stramenopiles, so this dataset provides the cell-biological baseline against which the derived photosynthetic ochrophytes and osmotrophic pseudofungi can be compared. It offers an investigative resource for exploring protein function, compartmental organization, and cellular architecture in a lineage whose ecology is far better understood than its cell biology. Investigators interested in using these data prior to publication are encouraged to contact the authors to avoid overlap and to facilitate coordinated and collaborative use of this resource.

## Data availability

The mass spectrometry proteomics data have been deposited to the ProteomeXchange Consortium via the PRIDE partner repository with the dataset identifier PXD085466 (Deutsch et al., 2020).

## Methods

### Culturing and Cell Harvesting

*Cafeteria roenbergensis* strain RCC4623 was cultured at room temperature in f/2 medium supplemented with 0.5 g/L tryptic soy broth (Millipore, 22092) and maintained as a mixed culture with an uncharacterized bacterial community serving as prey (Hackl et al., 2020). Cultures were grown to approximately ∼2.8 × 10^4^ cells/mL, as determined by haemocytometer counting. Cells were separated from residual bacteria and debris by filtration using 10 µm woven nylon mash filter (Tisch, ME17399), harvested by centrifugation at 6,000 × g for 10 min at room temperature, and washed twice, with centrifugation at 1,000 × g for 5 min after each wash. The final pellet was resuspended in ice-cold detergent-free (DF) buffer (0.25 M sucrose, 10 mM HEPES, pH 7.4, and 2 mM EDTA) supplemented with Halt™ Protease and Phosphatase Inhibitor Cocktail (Thermo Scientific), to a final volume of 8 mL.

### Cell Lysis and Fractionation

Cells were gently disrupted using three different methods: (i) nitrogen cavitation; (ii) passage through a stack of two 5.0 µm filter membranes; and (iii) sequential passage through 5.0 µm and 1.2 µm syringe filters. For nitrogen cavitation (i), the cell suspension was transferred into a cell disruption vessel (Parr 4639, Parr Instrument Co.) pre-chilled to 4 °C and connected to a nitrogen gas tank. The vessel was pressurized to 500 psi and equilibrated on ice for 5 min, followed by a reduction in pressure to 300 psi for an additional 5 min. After equilibration, the cell homogenate was collected through the outlet valve at approximately 3 drops/s, gently folding the foam into the liquid. For membrane filtration (ii), the cell suspension was passed through a stack of two 5.0 µm filter membranes (Millipore, SVLP02500) using a Swinnex Filter Holder (Millipore, SX0002500). For sequential syringe filtration (iii), the cell suspension was first passed through a 5.0 µm membrane syringe filter (Millipore, SLSV0R25LS) to remove cell aggregates, followed by a second passage through a 1.2 µm membrane syringe filter (Minisart, 17593). Subcellular fractionation was performed according to the differential centrifugation protocol described previously (Geladaki et al., 2019), generating 11 fractions: ten successive pellets and the supernatant remaining after the final spin. One set of fractions was reserved for assessment by SDS-PAGE, while the other was stored at –80 °C for downstream mass spectrometry. The experiment was performed in triplicates.

### SDS-PAGE and Coomassie Staining

Protein recovery and distribution across the fractionation series were assessed by SDS-PAGE. Pellets obtained from differential centrifugation were resuspended in 2× Laemmli buffer (4% SDS, 20% glycerol, 120 mM Tris-HCl pH 6.8) and heated at 95 °C until dissolved, and protein was quantified using the Pierce™ BCA Protein Assay Kit (Thermo Scientific). 10 µg of protein per fraction was separated on Bolt Bis-Tris Plus Mini Protein Gels, 4–12% (1.0 mm, WedgeWell format, Invitrogen) alongside protein marker (Amersham ECL Rainbow Marker, Full Range). Gels were stained for 1 h in 0.25% Coomassie brilliant blue (Pierce Coomassie Brilliant Blue R-250, Thermo Scientific) in methanol/water/acetic acid (1:10), destained by diffusion in the same solution minus the dye, and stored in ddH2O to clear the background. Gels were imaged using an Azure 600 (Biosystems) (Figure S1).

### Proteomics of Cafeteria Organellar Fractions by LC-MS/MS

Pelleted organellar fractions were resuspended in 500 µL lysis buffer (8 M urea, 75 mM NaCl, 1 mM EDTA, 50 mM Tris-HCl pH 8.0, 1× HALT Protease and Phosphatase Inhibitor Cocktail (Thermo Fisher Scientific)) and disrupted by sonication using four 15 s pulses at 50% amplitude, with 1 s intervals, for a total of 1 min. Insoluble material was removed by centrifugation at 20,000 rcf for 10 min at 4 °C, and total protein in the clarified supernatant was quantified by BCA assay (Thermo). Aliquots of 50 µg protein from each sample were reduced with 5 mM dithiothreitol (DTT), alkylated with 20 mM iodoacetamide (IAA), and digested overnight at 37 °C with Trypsin Gold (MS grade, Promega) at an enzyme:substrate ratio of 1:25. Peptides were purified by solid-phase extraction using 1 cc 50 mg C18 SPE columns (Waters Seppak), dried under vacuum, and stored at –80 °C. Dried peptides were resolubilized in 100 µL of 2% acetonitrile, 0.1% formic acid in water and their concentration reassessed by BCA assay. For spectral library construction, peptides from all samples were pooled in equal proportions and fractionated by high-pH reversed-phase HPLC on a C18 column (XBridge, 3.5 µm, 4.6 × 100 mm, Waters) at 30 °C and 0.5 mL/min over a 96 min gradient, yielding 96 fractions that were concatenated into 24 pooled fractions and dried under vacuum. Both individual samples and library fractions were resuspended in 2% acetonitrile, 0.1% formic acid in water supplemented with iRT peptides (Biognosys).

LC-MS/MS acquisition was performed on a Vanquish Neo ultra high-performance liquid chromatography (UHPLC) system (Thermo Fisher Scientific) interfaced with an Orbitrap Ascend Tribrid mass spectrometer (Thermo Fisher Scientific) fitted with a FAIMS Pro device. For each injection, 5 µL of sample was loaded directly onto a C18 analytical column (EasySpray ES802A, 75 µm inner diameter × 25 cm, 2 µm particle size, 100 Å pore size) held at 45 °C and eluted at 300 nL/min using a 120 min binary gradient of Solvent A (LC-MS grade water, 0.1% formic acid) and Solvent B (LC-MS grade acetonitrile, 0.1% formic acid). Spectral library fractions were measured in DDA mode with MS1 detection in the Orbitrap at 120K resolution, MS2 detection in the Orbitrap at 30K resolution, and HCD at an NCE of 25%. Individual samples were acquired in DIA mode with a scan range of 380–980 m/z, 16 m/z isolation windows with 1 m/z overlap, Orbitrap detection at 15K resolution, HCD NCE of 25%, and FAIMS CV of –45. A sample-specific spectral library was generated using the Pulsar search engine within Spectronaut (v19.0.240606.62635), searching against the *Cafeteria roenbergensis* strain RCC4623 predicted proteome (Hackl et al., 2020), with carbamidomethyl and oxidation as peptide modifications, up to two missed cleavages permitted, and proteins and peptides retained at a false discovery rate below 1%. No bacterial sequences were included in the search database.

DIA-MS data were then queried against the sample-specific spectral library in Spectronaut (v19.0.240606.62635) with default parameters, except that cross normalization was disabled. Protein and peptide identifications were accepted at a 1% FDR threshold. Abundance values exported from Spectronaut were normalized by variance stabilizing normalization in R (v4.3.0).

### Raw data processing

The resulting abundance profiles were imported into R (R Core Team, 2018) and analyzed using the Bioconductor packages MSnbase (v2.24.2; Gatto and Lilley, 2012) and pRoloc (v1.38.2; Gatto et al., 2014). The concatenated dataset from the 3 experiments yielded abundance profiles for 5,889 protein groups across 33 samples. The resulting table was imputed by neighbour averaging followed by zeroing. Principal component analysis and t-distributed stochastic neighbor embedding (t-SNE; van der Maaten and Hinton, 2008) were applied for dimensional reduction and data visualization.

### Supervised and Unsupervised Classification

A curated set of 571 marker proteins across 14 subcellular compartments was assembled (Table S1) and were used to train the classifier, which was then applied to all 5,889 protein groups. Supervised classification was performed with a support vector machine (SVM) model using the svmOptimization and svmClassification functions implemented in pRoloc (Gatto et al., 2014). For all clusters, the global median was used to set the cutoff score; proteins with an SVM score above the cutoff were considered predicted to their corresponding compartment, and those below were reset to “unknown”. The t-SNE map of the SVM-predicted proteins was plotted using the Shiny application implemented in pRolocGUI (v2.18.0).

In parallel, unsupervised k-means clustering was performed using the MLearn function from MLInterfaces (v1.88.1) in R to generate 14 clusters, within which protein distribution profiles are more similar to one another than to those of other clusters.

### Targeting sequence identification

TargetP 2.0 (Almagro Armenteros et al., 2019) was used to predict signal peptides (SP) and N-terminal mitochondrial targeting signals (MTS), with the organism group set to “Plant” to avoid database bias towards animal/fungi. DeepTMHMM v1.0.24 (Hallgren et al., 2022) was applied to predict transmembrane topology. Finally, nuclear localization signals (NLS) were identified using NLStradamus (Nguyen Ba et al., 2009) with the two-state static HMM and a posterior probability cutoff of 0.95.

## Acknowledgements

J.G.W. was supported by a grant from the National Science Foundation (DBI-2119963) and a grant from the Gordon and Betty Moore Foundation (GBMF10600). DJ was supported by VEDA FELLOWSHIPS within the Operational program Jan Amos Komensky (OP JAK), Project No. CZ.02.01.01/00/22_010/0008117. This research includes work conducted in the Integrated Mass Spectrometry Shared Resource supported by the National Cancer Institute of the NIH under grant number P30CA033572.

## Supporting Information

**Table S1.Marker proteins used in this study.**

**Table S2. SVM scores for all proteins detected in this study.**

**Figure S1.**
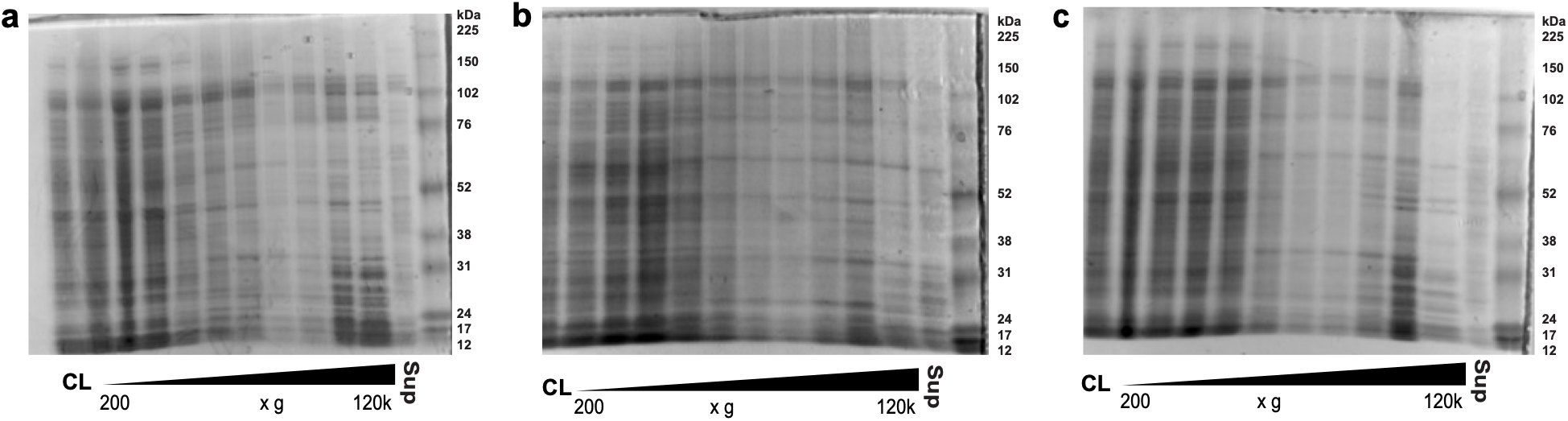
Coomassie-stained SDS-PAGE of subcellular fractions of *Cafeteria roenbergensis*. Fractions are arranged in order of increasing centrifugation speed, with cell lysate and the post-ultracentrifugation supernatant in the first and last lanes. We used three different methods for cell lyses (A) nitrogen cavitation (500 psi/5 min, followed by 300 psi/5 min); (B) passing through a stack of two 5.0 µm filters; and (C) passage through a single 5 µm membrane filter and then through 1.2 µm membrane syringe filter. Differences in band pattern across the series indicate that protein was partitioned between fractions rather than carried through the series uniformly.

**Figure S2.**
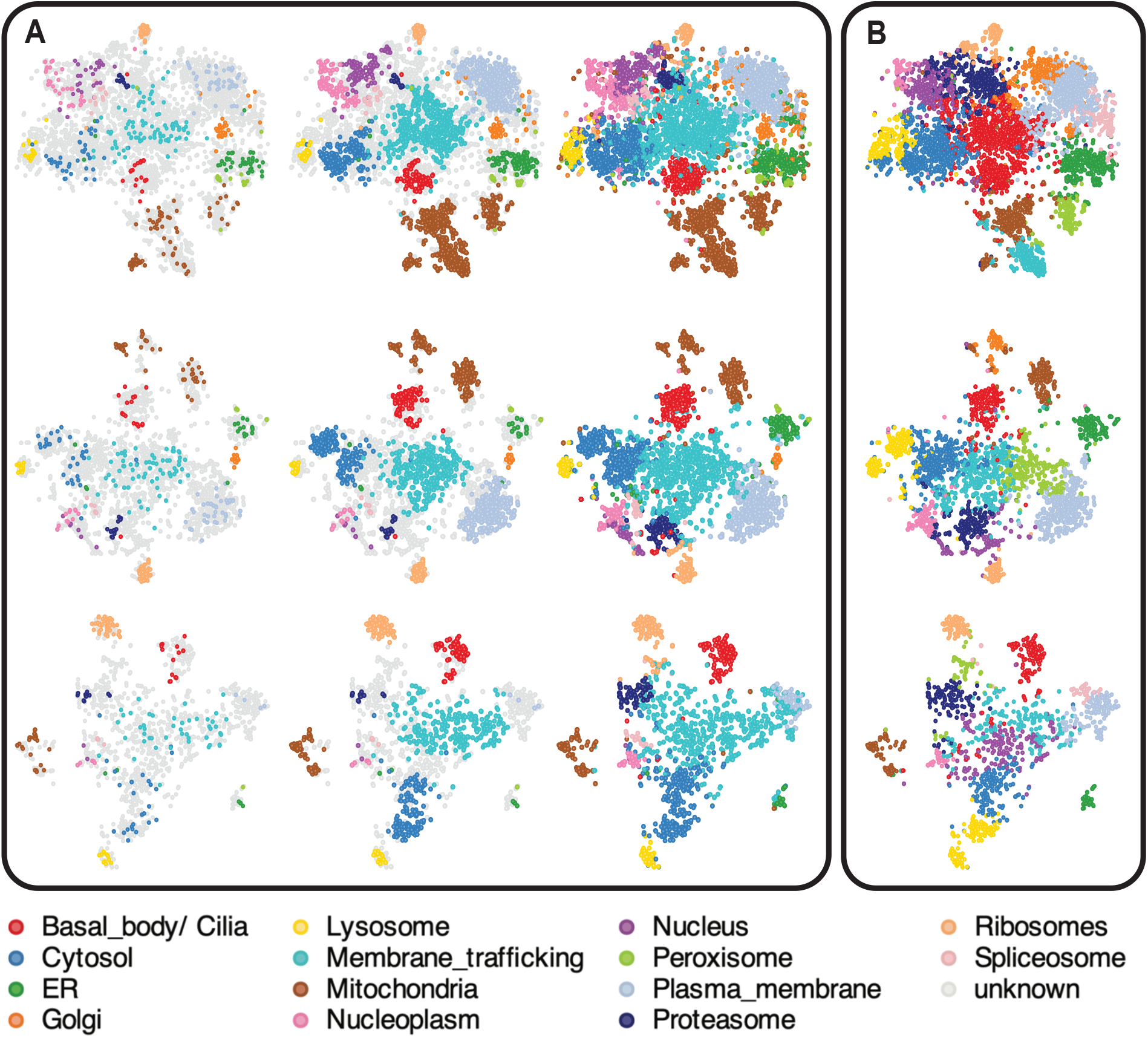
Subcellular atlases of *Cafeteria roenbergensis* under different imputation strategies and k-means clustering. A. SVM classification of datasets with neighbour averaging followed by zeroing (top row), neighbour averaging (middle row), and with no imputation (bottom row). For each, the marker set, the assignments retained at the median SVM cutoff, and all assignments with no cutoff are shown from left to right. B. Unsupervised k-means clustering with k = 14, for the same three imputation strategies. k-means clusters carry no organelle identity of their own; each cluster has been given the colour of the marker-defined compartment it most closely corresponds to, so that agreement between the unsupervised and supervised classifications can be read directly from the figure.

